# Mycoplasma Faucium and Breast Cancer

**DOI:** 10.1101/089128

**Authors:** V. Mitin, L. Tumanova, N. Botnariuc

## Abstract

Viruses and bacteria are the cause of a large number of different human diseases. It is believed that some of them may even contribute to the development of cancer. The present work is dedicated to the identification of mycoplasmas in patients with breast cancer. Mycoplasmas may participate in the development of several human diseases including chronic fatigue syndrome, acquired immunodeficiency syndrome, atypical pneumonia, etc. Moreover, there is a reason to believe that mycoplasma can participate in the development of cancer, leukemia and lymphoma.

DNA samples from blood, saliva and tumor tissues of the Oncology Institute of Moldova patients diagnosed with breast cancer were analyzed. Mycoplasma testing was performed using nested PCR method. For *Mycoplasma spp*. detection, we used primers from the region of the 16S-23S RNA genes. The identification of *Mycoplasma faucium*, *Mycoplasma salivarium* and *Mycoplasma orale* was performed by nested PCR with primers for RNA polymerase beta subunit gene corresponding to mycoplasma.

*M.faucium* and *M.salivarius* was found in saliva at about 100%, and *M.or*ale at a frequency of about 50%. Only *M.faucium* was found with the frequency of about 60% in the tissue of the patients. Moreover, a fairly high rate of detection of mycoplasma is observed both in the cases when primers for RNA polymerase gene and primers for 16S-23S RNA were used.

We found *M.faucium* in tumor tissues of patients diagnosed with breast cancer. It is known that mycoplasmas are able to stimulate the synthesis of certain cytokines, which act as mitogenes on the cell. We assume that M.faucium can stimulate the mitogenes synthesis in breast tissues (e.g., cytokines) which, in turn, stimulate cell division and thus participate in the initiation of breast cancer.

## Background

Viruses and bacteria are the cause of a large number of different human diseases. It is believed that some of them may even contribute to the development of cancer. We assume that the bacterial and viral infections may be one of the causes of breast cancer. This assumption is evidenced by published data showing that in the tumor tissue of breast cancer patients *Epstein-Barr* virus (EBV) and human mammary tumor virus (HMTV) are often detected [1].

We have analyzed the possibility to detect a number of viruses and bacteria in tissues from patients with breast cancer. Preliminary tissue analysis of 10 patients with breast cancer has been carried out by PCR. *Epstein-Barr* virus, *Cytomegalovirus, Herpes simplex* virus, *Herpes* viruses of types *6, 7, 8*, as well as as bacteria-*Chlamydia trachomatis, Mycoplasma hominis, Mycoplasma genitalium, Mycoplasma complex* (*Mycoplasma complex* set, detects a group of mycoplasmas), *Ureaplasma urealiticum* and *Gardnerella vaginalis* ware analyzed. Only *Epstein-Barr virus*, *Gardnerella vaginalis* and *Mycoplasma complex* were detected in the tissues after analyzing all these viruses and bacteria. These pathogens were analyzed in a more detailed study. The results of analysis of *Gardnerella vaginalis* were previously published [2].

The present work is dedicated to the identification of mycoplasmas in patients with breast cancer. Mycoplasma is a genus of bacteria, which is formed by a large group of microorganisms different from the conventional bacteria by a small size and the lack of cell wall. Lack of a rigid cell wall allows a close contact with the cytoplasmic membrane of Mycoplasma cell. Under these conditions, mycoplasma is hardly recognized by the immune system as a foreign organism. In addition, mycoplasma is capable of penetrating into the cell and in this way becomes an intracellular parasite. The intracellular location is a niche in which the mycoplasma is well protected from the immune system and the effect of many antibiotics. Mycoplasmas have a very small genome and, therefore, their survival depends on the host. Mycoplasmas have to use proteins, nucleic acids, cholesterol, fatty acids, etc., received from the host cells, to survive. At the same time, a disproportionately large number of mycoplasma genes are involved in cell adhesion problem and genes that contribute to the maintenance and care of parasitism surveillance by the immune system (for example, genes responsible for the synthesis of membrane antigens). Apparently, one of the main directions of mycoplasma’s evolution in resolving the problem of survival is to become invisible to the immune system of the host organism. Mycoplasmas may participate in the development of several human diseases including chronic fatigue syndrome, acquired immunodeficiency syndrome, psoriasis, scleroderma, atypical pneumonia, cystitis, and Alzheimer's disease, etc. Moreover, there is a reason to believe that mycoplasma can participate in the development of cancer, leukemia and lymphoma [3].

## Methods

DNA samples from blood, saliva and tumor tissues of the Oncology Institute of Moldova patients diagnosed with breast cancer were analyzed. Blood and saliva collection from breast cancer patients performed in the Institute of Oncology of Moldova 1 – 2 hours before tumor resection, and tissue samples – after tumor resection. All samples were stored and transported at +4°C. Time between sample collection and DNA isolation did not exceed 24 hours.

DNA from blood and saliva and tumor tissues was isolated using lysis buffer containing 5M guanidinium thiocyanate, 50mM Tris (pH 7.5), 10mM EDTA, 0.5% Triton X-100, then extracted with equal volume of phenol:chlorophorm and then chlorophorm. DNA was precipitated by 2.5 volumes of ice-cold 96% ethanol in the presence of 0.3M ammonium acetate. Tumor was grinded in liquid nitrogen with addition of lysis buffer before extraction.

Mycoplasma testing was performed using nested PCR method. For *Mycoplasma spp*. detection, we used primers from the region of the 16S-23S RNA genes and primers for the RNA polymerase beta subunit gene (Table 1).

**Table 1.**
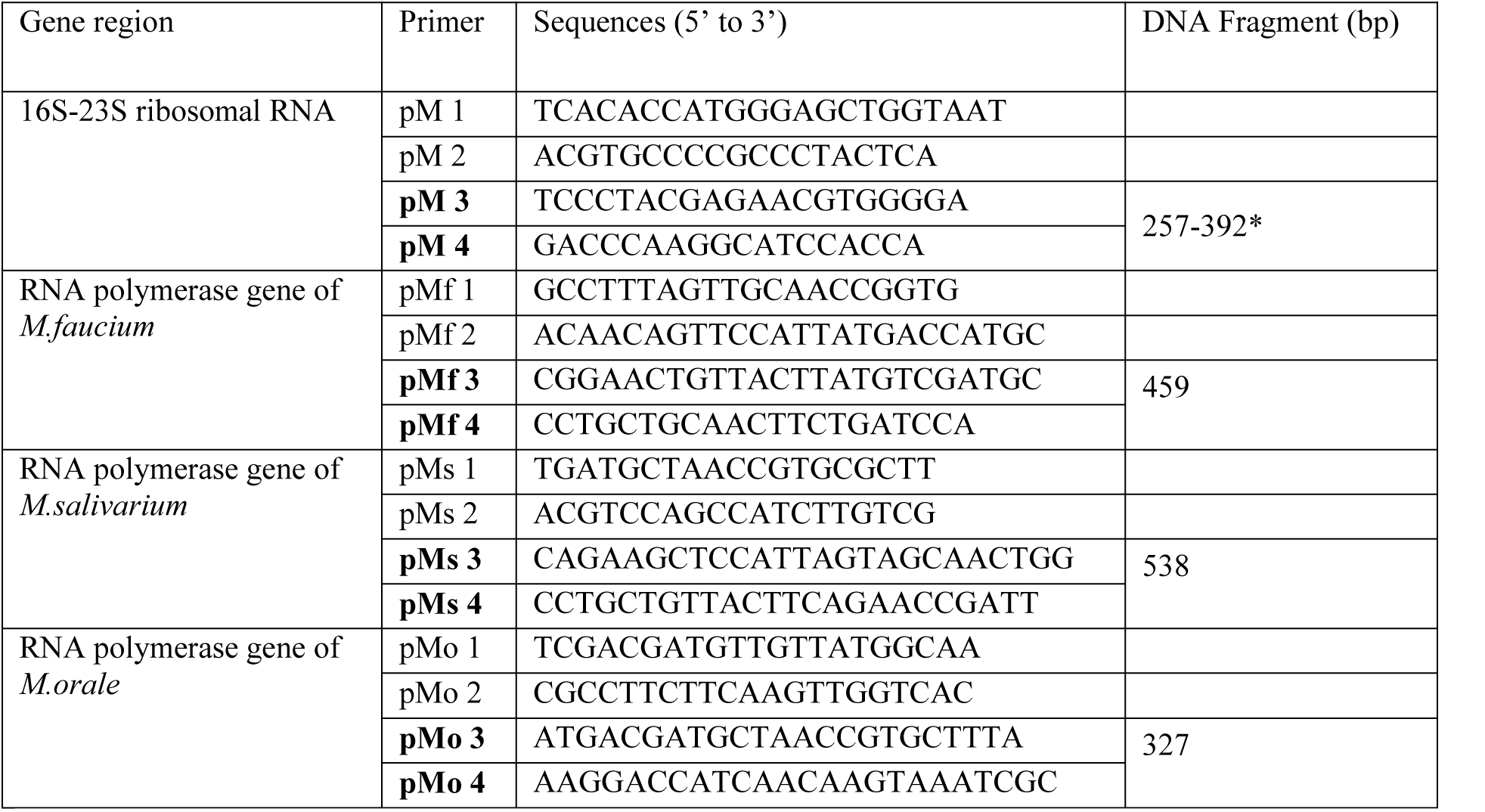
Primer sequences for *Mycoplasma spp*. testing

*range of DNA fragment sizes for different mycoplasmas is mentioned in the Results section. In bold are the inner primers for nested PCR analysis.

Primers were synthesized by Alpha DNA (Montreal, Quebec, Canada).

Both rounds were 30 cycles, with 60°C annealing temperature. Each of the samples was analyzed twice. The amplification products were analyzed in 2% agarose gel. 100 bp. ladder (Fermentas) was used as a molecular marker.

### Results and discussion

Identification of *Mycoplasma spp*. in patients with breast cancer was carried out using primers from the region of the 16S-23S RNA genes. This region includes 16S ribosomal RNA gene, partial sequence; 16S-23S ribosomal RNA intergenic spacer, complete sequence; and 23S ribosomal RNA gene, partial sequence. The length of intergenic region for any mycoplasma varies depending on the type of mycoplasma. So, use of 16S-23S primers allows not only to detect the mycoplasma, but also to effectuate a preliminary analysis of mycoplasma species. A number of human mycoplasmas, such as -*M.buccale* (296 bp), *M.faucium* (286 bp), *M.fermentans* (387 bp), *M.hominis* (257 bp), *M.lipophilum* (373 bp), *M.orale* (311 bp), *M.primatum* (388 bp), *M.salivarium* (290 bp), *M.spermatophilum* (392 bp) may be identified using these primers.

Based on the length of DNA fragments, synthesized by nested PCR with 16S −23S primers (~ 280 bp. in the course of tumor tissue analyzing) we selected *M.faucium*, *M.orale* and *M.salivarium* for further analysis. In this, more detailed analysis we used primers for RNA polymerase beta subunit gene. Figure 1A shows the result of analysis of DNA samples for blood, saliva and tumor tissue of four patients using 16S −23S primers. Figures 1B, 1C and 1D show results of analysis of saliva samples with primers, specific for RNA polymerase gene.

**Figure 1.**
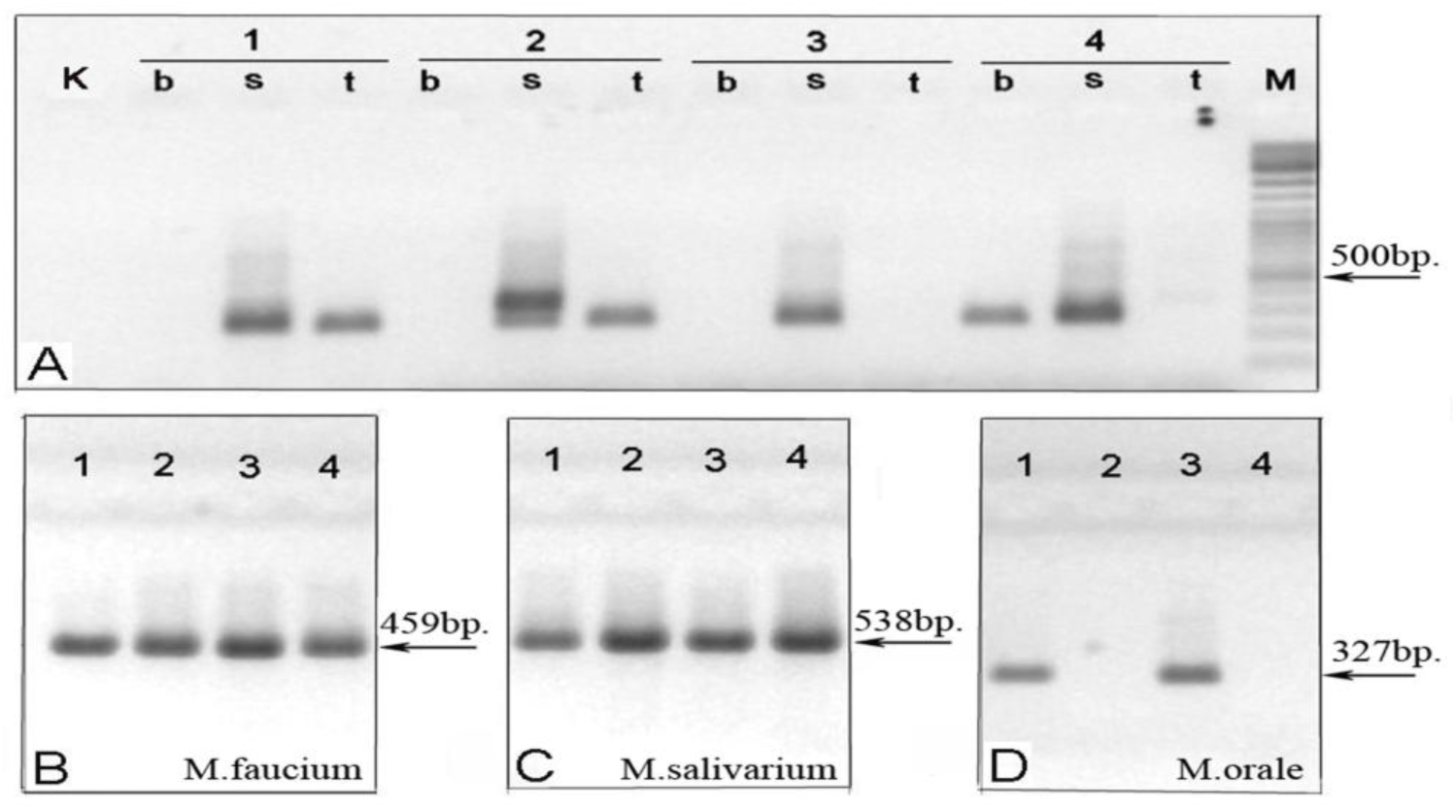
Analysis of the nested PCR obtained with the mycoplasma-specific primers on DNA of four (1, 2, 3, 4) patients. (A) - PCR analysis of DNA from the blood (b), saliva (s) and tumor tissues (t) with primers from the region 16S-23S RNA genes. (K) - negative control; (M) - molecular marker. (B, C, D) - the detection of *M. faucium*, *M. salivarius* and *M.orale* in saliva using primers from RNA polymerase gene.

The length of the synthesized DNA fragments using the 16S-23S primers varied in the course of saliva samples analysis and was constant in the tumor tissues analysis. This can be explained by the fact that the saliva samples usually contain several types of mycoplasma. In this case electrophoregram pattern depends not only on the types of mycoplasma but also on the ratio of their concentrations. Constant fragment size during tumor tissues analysis may indicate that only one type of mycoplasma was present. The specificity was experimentally confirmed by PCR analysis *M.faucium* in the presence or absence *M.salivarium*, most similar to *M.faucium* mycoplasma according to nucleotide sequence analysis. In addition, alignment of RNA polymerase beta subunit genes of the nine listed above mycoplasmas also points to the specificity of each pair of primers used in this work (results are not shown).

Table 2 shows the results of detection of various mycoplasmas in tissue, blood and saliva of breast cancer patients. It is observed that only *M.faucium* may be found in the tissue of patients. Moreover, a fairly high rate of detection of Mycoplasma *M.faucium* is observed both in the cases when specific primers (for RNA polymerase gene) and primers for 16S-23S RNA, which are able to detect various mycoplasmas, including *M.faucium* were used.

**Table 2.** Identification of mycoplasmas in biological samples of patients diagnosed with breast cancer.

| Biological sample | 16S-23S |  | *M.faucium* |  | *M.salivarium* |  | *M. orale* |  |
| --- | --- | --- | --- | --- | --- | --- | --- | --- |
|  | % positive | Total number | % positive | Total number | % positive | Total number | % positive | Total number |
| Tumor tissue | 42.86 | 28 | 60.7 | 28 | 0 | 15 | 0 | 15 |
| Blood | 19.3 | 57 | 9.4 | 32 | 0 | 21 | 4.8 | 21 |
| Saliva | 96.9 | 65 | 100 | 31 | 100 | 20 | 55 | 20 |

The same time, in cases where DNA from saliva was analyzed, approximately 100% of the samples were positive for both *M.faucium* and *M.salivaium*. Low frequency detection of mycoplasma in blood, together with the high frequency of detection of mycoplasma in tissue and saliva indicates the absence of false positive diagnosis of *M.faucium*.

Earlier [4], in breast cancer patients *M.hyorhinis* has been detected. Nevertheless, we failed to confirm this conclusion, using the primers of the p29 gene, specific for *M.hyorhinis*, (for 10 patients, data not shown). This may be due to the different localization of the Chinese and Moldavian groups of patients. The results of these studies argue high frequency of *M.faucium* detection in tumor tissue of breast cancer patients. But does this mean that *M.faucium* can really participate in the development of breast cancer? What can be said in favor of such assumption? Unfortunately, almost nothing is known about molecular biology of *M.faucium*, but its properties may be similar to those of other mycoplasmas. For example it is known that mycoplasmas trigger the secretion of cytokines, including IL-1b (interleukin-1b), IL-2, IL-4, IL-6, IL-8, IL-10, interleukin-2 receptor (IL-2R), interferon alpha (IFN-α), interferon gamma (IFN-γ), tumor necrosis factor alpha (TNF-α), and granulocyte macrophage-colony stimulating factor (GM-CSF)[5]. However, most of these cytokines are growth factors. This means that when they interact with appropriate receptors, D cyclins and cyclin-dependent kinases 4/6 (cdk 4/6) appear in the cell, which in turn initiate cell division. If the growth factor receptors located on the cell membrane, which are present in breast tissue, it may mean that such cytokines may promote cell division in a mammary gland. At the same time, cell division which is not controlled by the organism is one of the main (if not the main) symptom of cancer. Thereby mycoplasmas theoretically are able to participate in the development of cancer by activating cell division. Data presented in the Table 3 show that some cytokines can participate in the development of cancer.

**Table 3.** Some cytokines (growth factors) may participate in a cancer development

| Cytokines * | Cytokines receptors (CD markers) | Cytokines receptors in breast tissues | Regulation of genes expression (cytokines as mitogenes) | Cytokines and cancer development |
| --- | --- | --- | --- | --- |
| IL-1β | IL-1R2 (CD 121) | fibroblasts, endothelial cells | Akt[6, 7], beta-catenin [8] | Colon cancer [9], breast cancer [10]. |
| IL-2 | IL-2R (CD122) |  | Akt [11], Stat5 [12] | cervical cancer [13, 14] |
| IL-4 | IL-4R (CD124) | fibroblasts, endothelial cells | MAPK [15], Stat6 [16] | prostate cancer [17, 18] |
| IL-6 | IL-6R (CD126) |  | STAT3 [19], RAS [20]. | breast cancer [21, 22] |
| TNF-α | FAS (CD95) | epithelium and connective tissue | MAPK, JNK, Akt [23], MAPK [24] | fibroblast proliferation[25], breast tumor [26] |
| GM-CSF | GM-CSFRα (CD116) | fibroblasts, endothelial cells | cyclin D1[27]. | lung cancer [28, 29]. |

\* IL-1β - activates T cells. IL-2 - T-cell growth factor, stimulates IL-1 synthesis, activates B-cells and NK cells. IL-4 - growth factor for activated B cells, resting T cells, and mast cells. IL-6 - stimulates Ig synthesis, growth factor for plasma cells. TNF-α - a member of the epidermal growth factor (EGF) family, TGF-α is a mitogenic polypeptide. GM-CSF - it is a white blood cell growth factor.

* IL-1β-activates T cells. IL-2- T-cell growth factor, stimulates IL-1 synthesis, activates B-cells and NK cells. IL-4-growth factor for activated B cells, resting T cells, and mast cells. IL-6-stimulates Ig synthesis, growth factor for plasma cells. TNF-a-a member of the epidermal growth factor (EGF) family, TGF-α is a mitogenic polypeptide. GM-CSF-it is a white blood cell growth factor.

But these cytokines secretion may be triggered by mycoplasmas. This means that the mycoplasma can initiate division of those cells, which contain corresponding cytokine receptors. Since fibroblasts and endothelial cells contain receptors for above-mentioned cytokines it can be expected that mycoplasmas are able to participate in the development of breast cancer.

## Conclusion

We detected *M.faucium* in tumor tissues of patients diagnosed with breast cancer. It may be assumed that this particular mycoplasma is somehow involved in breast cancer development.However, the detection of it in tumor tissues of patients diagnosed with breast cancer, and the data that *Mycoplasma spp*. could theoretically cause cell division, is unlikely to be enough to claim that *M.faucium* is involved in the cancer development. A stronger evidence for this would be increasing of the efficacy of breast cancer therapy by eradication of *M.faucium*.

